# Local CpG-*Stat3* siRNA treatment improves antitumor effects of immune checkpoint inhibitors

**DOI:** 10.1101/2023.08.17.553571

**Authors:** Chunyan Zhang, Rui Huang, Lyuzhi Ren, JiEun Song, Marcin Kortylewski, Piotr Swiderski, Stephen Forman, Hua Yu

## Abstract

Immune checkpoint blockade (ICB) therapy has significantly benefited patients with several types of solid tumors and some lymphomas. However, many of the treated patients do not have durable clinical response. It has been demonstrated that rescuing exhausted CD8^+^ T cells is required for ICB-mediated antitumor effects. We recently developed an immunostimulatory strategy based on silencing STAT3 while stimulating immune responses by CpG, ligand for Toll-like receptor 9 (TLR9). The CpG-small interfering RNA (siRNA) conjugates efficiently enter immune cells, silencing STAT3 and activating innate immunity to enhance T-cell mediated antitumor immune responses. In the present study, we demonstrate that blocking STAT3 through locally delivered CpG-*Stat3* siRNA enhances the efficacies of the systemic PD-1 and CTLA4 blockade against mouse A20 B cell lymphoma. In addition, locally delivered CpG-*Stat3* siRNA combined with systemic administration of PD-1 antibody significantly augmented both local and systemic antitumor effects against mouse B16 melanoma tumors, with enhanced tumor-associated T cell activation. Overall, our studies in both B cell lymphoma and melanoma mouse models demonstrate the potential of combinatory immunotherapy with CpG-*Stat3* siRNA and checkpoint inhibitors as a therapeutic strategy for B cell lymphoma and melanoma.

## INTRODUCTION

Therapeutic immune checkpoint blockade (ICB) has profoundly and positively impacted B cell lymphoma and melanoma treatment. However, many of treated patients do not respond and those who have good initial response may not have durable clinical benefits^1–3^. The limited responses to ICB in cancer patients are largely attributed to lack of IFN signaling and activation of CD8^+^ T cells even after immune checkpoint is removed^4–7^. Extensive studies from our group and others have demonstrated that IFNγ, which is necessary for PD-1 directed therapy responses^8, 9^, is inhibited in tumor-associated immune cells by STAT3^10^. STAT3 is also known to promote tumor cell proliferation, survival, and invasion in diverse cancers^11–16^. The immunosuppressive role of STAT3 in tumor cells, T cells, B cells, myeloid cells and dendritic cells has been documented extensively^10–23^. Additionally, STAT3 signaling in CD4^+^ T cells promotes Treg accumulation in tumors, while inhibiting CD8^+^ T_EFF_ tumor infiltration and antitumor immunity^12, 18^.

STAT3 as a target for inhibiting B cell lymphoma cell survival and activating antitumor immune response has been documented^24–27^. We previously developed an immunostimulatory strategy by linking *Stat3* siRNA with CpG, the ligand for TLR9^20, 25, 28–30^. CpG not only facilitates the delivery of the siRNA but, in the absence of STAT3 activity, it also activates potent antitumor immune responses^30^. We further showed that both CpG-*Stat3* siRNA and CTLA4 antibody inhibit STAT3 in B malignant cells, leading to tumor cell apoptosis and/or proliferation inhibition^25, 31^. Additionally, CpG-*Stat3* siRNA activates antitumor T cells through blocking STAT3 in macrophages, B cells and DCs while CTLA4 or PD-1 antibody also enables interaction between DCs and T cells to activate antitumor T cell immunity^20, 32, 33^. Thus, combinatory treatments with CpG-*Stat3* siRNA and CTLA4 or PD-1 antibody may significantly boost the antitumor efficacies of CTLA4 or PD-1 antibody in treating B cell lymphoma and melanoma.

Local treatment can lead to direct anti-tumor effects at the injected tumor sites and induction of systemic anti-tumor immune responses^34–37^. Both lymph node-resident B cell lymphoma and the cutaneous and subcutaneous melanoma tumors provide the opportunity for local treatment for easier delivery and generally lower toxicity. Previously, we showed that local treatments with CpG-*Stat3* siRNA inhibit both B cell lymphoma^25^ and melanoma tumor growth^20^, while resulting in effective tumor growth inhibition including complete tumor eradication when combined with localized radiation therapy^25, 38^. However, whether CpG-*Stat3* siRNA can boost the efficacies of ICB in these two tumor types remains unknown. The CpG oligonucleotide TLR9 ODN 1826 agonist has been shown to effectively augment the therapeutic potential of checkpoint blockade through locally delivered ODN 1826 and systemic CTLA4 or PD-1 blockade antibody in the B16 melanoma mouse model^37^. However, high CpG dosing (30 μg/intratumorally) was required to enhance ICB response in this study^37^. High dose CpG injections induced a sharp and sustained increased local and serum levels of IL-12, IL-6, and TNF-α, followed by high levels of the acute-phase proteins serum amyloid A (SAA) and serum amyloid P (SAP), which could cause strong side effects^39, 40^. Therefore, we hypothesized that CpG-*Stat3* siRNA, at significantly lower and safer effective CpG concentrations, may enhance anti-tumor efficacy of ICB by blocking STAT3 in the tumor-associated myeloid cells, thereby stimulating antigen-presentation and activating CD4^+^ and CD8^+^ T cell-mediated antitumor immune responses. Our study shows that at relatively low CpG-*Stat3* siRNA concentration, local injection can augment the therapeutic potential of PD-1 antibody not only in CpG-*Stat3* siRNA treated tumors but also in distal non-treated tumors.

## RESULTS

### Local CpG-*STAT3* siRNA treatment enhances anti-lymphoma effects of CTLA4 and PD-1 blockade

Although some advances have been made in treating cancer patients with ICB, the efficacy is limited, which is contributed by the lack of CD8^+^ T cell activation^2, 6^. We have shown previously that CpG-*Stat3* siRNA treatment in vivo silences *Stat3* in dendritic cells, B cells and macrophages, leading to T cell activation and potent antitumor immunity^20^. We further showed that both CpG-*Stat3* siRNA and CTLA4 antibody inhibit STAT3 in B malignant cells, leading to tumor cell apoptosis and/or growth inhibition^25, 31^. Here we assessed whether silencing *Stat3* with CpG-*Stat3* siRNA can significantly enhance the efficacies of CTLA4 antibody therapy in B cell lymphoma by augmenting the immunostimulatory effects. For our experiments, we selected commonly used mouse A20 B cell lymphoma model, which exhibits TLR9 expression and constitutive Stat3 activation^25^. We have previously shown that CpG *Stat3* siRNA efficiently targeted A20 tumor cells to silence *Stat3*^20, 25^. To evaluate gene silencing effect CpG-*Stat3* siRNA, we used quantitative real-time PCR (qPCR) analysis of *Stat3* mRNA of CpG-*Stat3* siRNA treated A20 tumor cells. The effect of *Stat3* gene silencing was observed in cultured A20 cells at 48 h since the start of CpG-*Stat3* siRNA treatment **(Figure 1A).** Next, we tested the feasibility of using this strategy for targeting Stat3 in vivo. BALB/c mice with established, subcutaneously engrafted (s.c.-engrafted) A20 lymphoma tumors were treated with PBS, CpG-*Stat3* siRNA at 12.5 μg or 25 μg/mouse by daily intra-tumoral (IT) injections. One day after the third injection of PBS or CpG-*Stat3* siRNA, mice were euthanized, and the tumor tissues were harvested to assess *Stat3* mRNA expression by qPCR. The CpG-*Stat3* siRNA treatment reduced the *Stat3* mRNA expression in a dose-dependent manner **(Figure 1B).**

**Figure 1.**
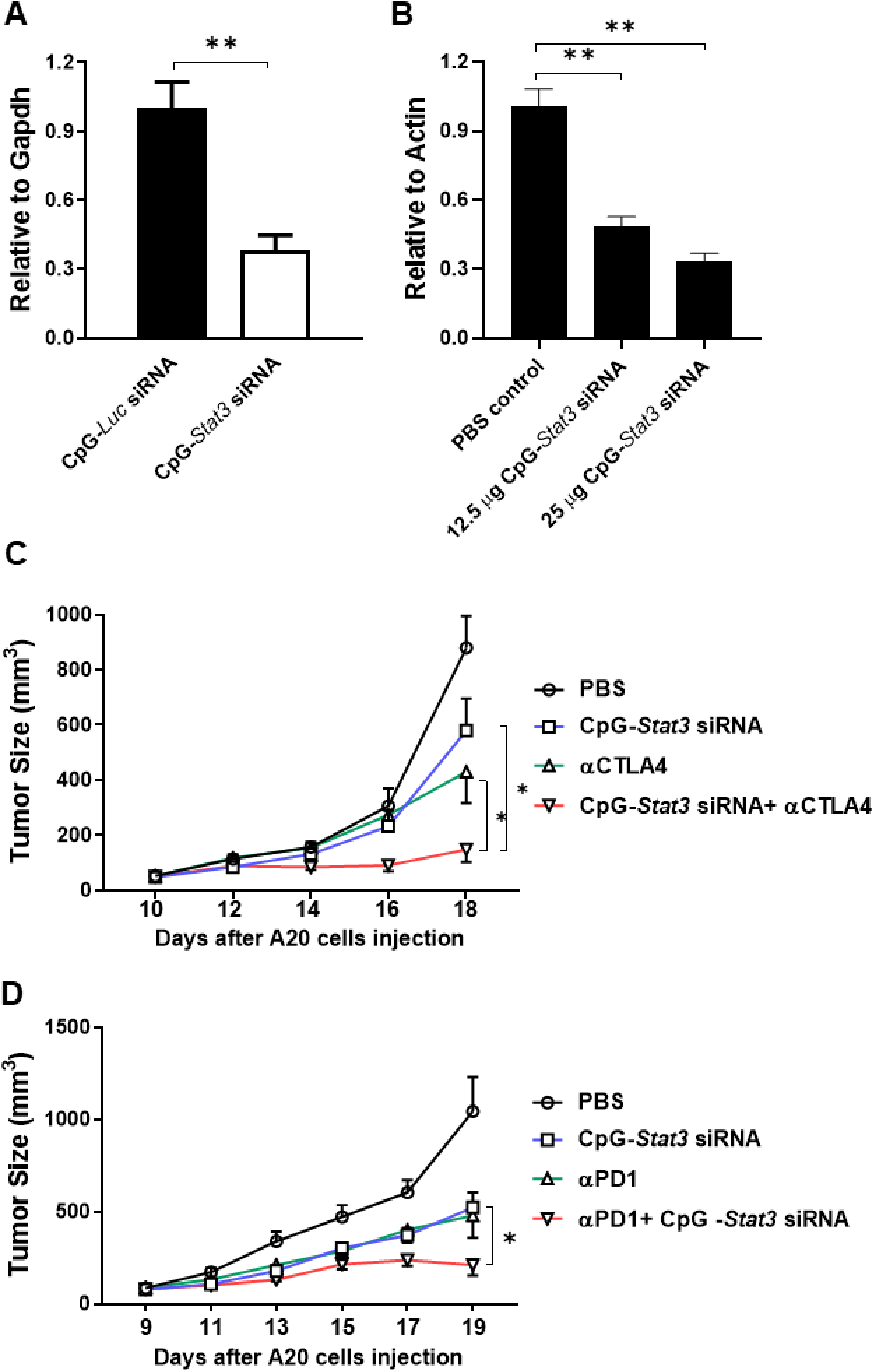
CpG-*Stat3* siRNA gene silencing effect in A20 B cell lymphoma cells and therapeutic effect of CpG-*Stat3* siRNA with or without CTLA4 or PD-1 blockade. (A) A20 cells were cultured in the presence of CpG-*Stat3* siRNA or CpG-*Luc*-siRNA for 48 h. CpG-*Stat3* siRNA silences *Stat3* gene expression at mRNA level is shown by q-PCR. Data are shown as means ± SEM, n = 3. (B) BALB/c mice with subcutaneous (s.c) A20 lymphoma tumors were treated by intra-tumoral injections of CpG-*Stat3* siRNA, using indicated amounts or PBS. CpG-*Stat3* siRNA gene silencing effect is shown by q-PCR. Data are shown as means ± SEM, n = 3 mice. (C) BALB/c mice with s.c. A20 lymphoma tumors were treated by intra-tumoral injections of CpG-*Stat3* siRNA, i.p injection of anti-CTLA4, anti-PD-1 antibodies or combination treatment with CpG-*Stat3* siRNA and anti-CTLA4 or anti-PD-1 antibodies every other day, starting 9-10 day after tumor implantation (5 x 10^5^ A20 cells/tumor). Data are shown as means ± SEM, n=5. Student’s *t*-test was used for statistical analysis (\**P*<0.05; \*\**P*<0.01; \*\*\**P*<0.001). Tumor size was monitored every other day.

We next tested whether combining CpG-*Stat3* siRNA with CTLA4 or PD-1 inhibition would lead to superior antitumor effects than the single agents in the mouse B cell lymphoma. As shown in **Figure 1C and 1D**, silencing *Stat3* through locally delivered CpG-*Stat3* siRNA significantly enhanced the antitumor efficacies of systemic administration of PD-1– and CTLA4-specific antibodies in mice bearing the A20 B cell lymphoma arresting tumor growth.

### Combining CpG-*Stat3* siRNA with CTLA4 or PD-1 blockade induces significantly higher activities of tumor infiltrating T cells

Tumor infiltration by activated and not exhausted, cytotoxic CD8^+^ T cells at minimal presence of regulatory T lymphocytes is critical for successful outcome of ICB therapy^2, 6, 41^. Therefore, we determined whether T cell activation is a critical contributor for the antitumor effect of combinatory immunotherapy with CpG-*Stat3* siRNA and checkpoint blockades. Combinatory treatment with CpG-*Stat3* siRNA and CTLA4 antibody in A20 lymphoma tumor bearing mice significant increased the percentages of IFNγ and/or granzyme B (GZMB) producing CD4^+^ T cells and CD8^+^ T cells in tumors, compared to either CpG-*Stat3* siRNA or CTLA4 antibody alone **(Figure 2A-C)**. Similarly, combinatory treatments with CpG-*Stat3* siRNA and PD-1 antibody in A20 lymphoma tumor bearing mice led to significantly higher percentages of activated CD8^+^ T cells positive for IFNγ, CD107α, which measures T cell cytotoxicity, and GZMB at the tumor sites **(Figure 2D-F).** Furthermore, combinatory treatment led to inhibition of FoxP3^+^ regulatory T cells **(Figure 2G)**. These results support that local CpG-*Stat3* siRNA treatment enhances anti-tumor effects of CTLA4 or PD-1 blockade through increasing the antitumor effector functions of tumor infiltrating T cells.

**Figure 2.**
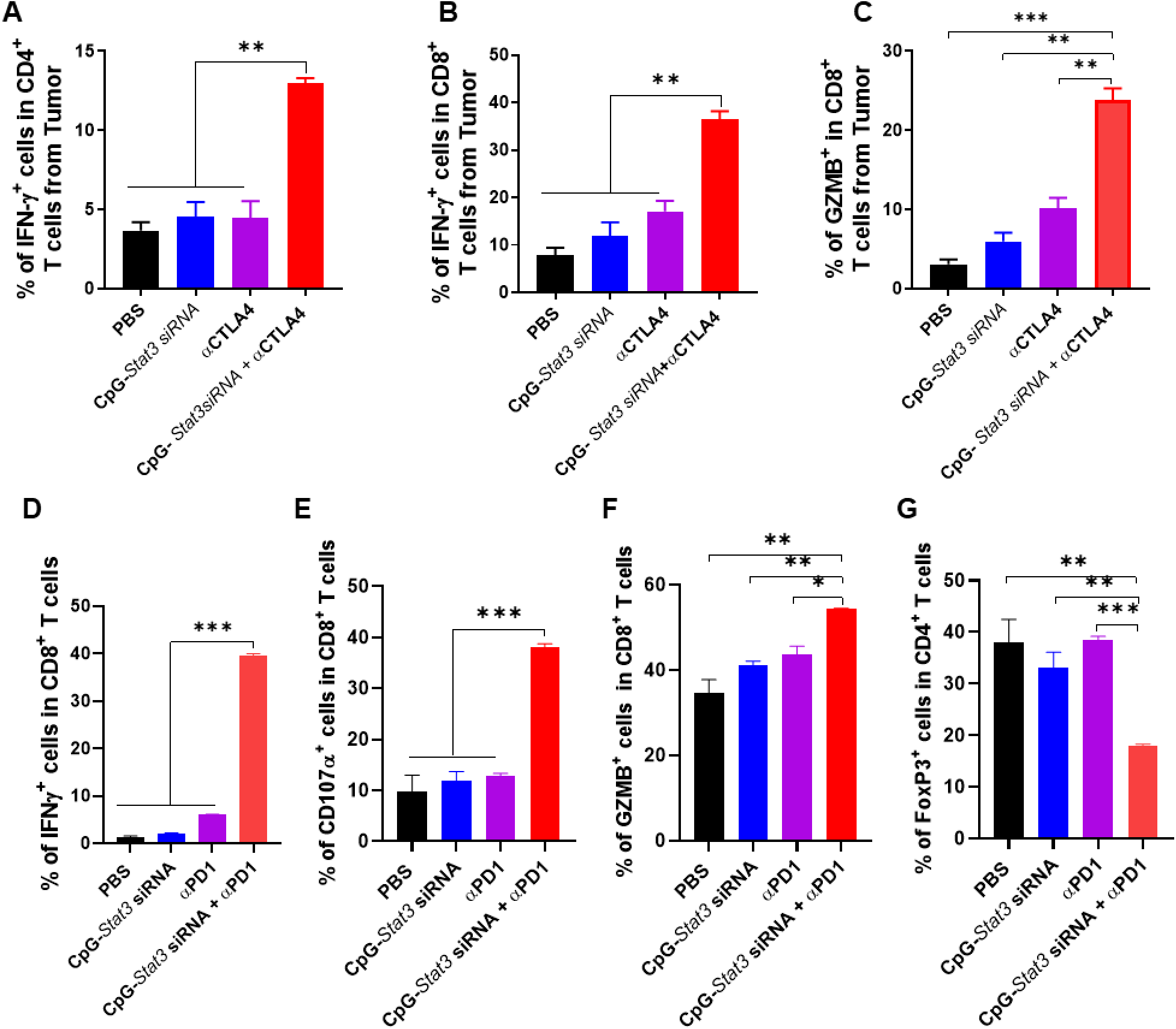
Local CpG-*Stat3* siRNA treatment significantly enhances the antitumor effector functions of tumor infiltrating T cells in mice treated with CTLA4 or PD-1 blockade. Single cell suspensions prepared from A20 tumors from mice receiving indicated treatments were analyzed by flow cytometry for IFNγ (A, B and D), CD107α (E), granzyme B (C and F) and FoxP3 (G) expression in T cells. Flow cytometry data showing IFNγ^+^ or GzmB^+^ cell frequencies in tumor-infiltrating CD4^+^ or CD8^+^ T cells. Data are shown as means ± SEM (n=3, each sample was pooled from 2-3 mice), Student’s *t*-test was used for statistical analysis (*P<0.05; **P<0.01; ***P<0.001).

### Local CpG-*STAT3* siRNA combined with PD-1 antibody results in systemic antitumor effects against melanoma in mice

We previously showed that local treatments with CpG-*Stat3* siRNA inhibit melanoma tumor growth by silencing *Stat3* and activating tumor infiltrating immune cells^20^. However, the ideal cancer therapy should not only inhibit local tumor regression but also induce a systemic anti-tumor immunity that could effectively eradicate distant metastases. Whether CpG-*Stat3* siRNA can enhance both local and systemic anti-tumor response of ICB in melanoma remains unknown. Therefore, we evaluated whether combinatory treatment with locally delivered CpG-*Stat3* siRNA and systemic PD-1 antibody treatment could lead to both local and systemic antitumor effects. We used a bilateral tumor model in which the C57BL/6 mice were challenged at both flanks by mouse B16 melanoma tumor cells through s.c. injection. Only one tumor was treated with locally delivered CpG-*Stat3* siRNA to assess local and systemic effects of single and combinatory treatments. Our results show that local CpG-*Stat3* siRNA treatment suppressed the growth of both treated and non-treated distal tumors **(Figure 3),** suggesting the generation of systemic antitumor immune responses. In addition, combination treatment with CpG-*Stat3* siRNA and systemic PD-1 antibody significantly enhanced the systemic antitumor responses compared to either CpG-*Stat3* siRNA or PD-1 antibody treatment alone **(Figure 3)**. These findings provide evidence that CpG-*Stat3* siRNA local treatment can enhance the PD-1 antibody mediated systemic anti-tumor response.

**Figure 3.**
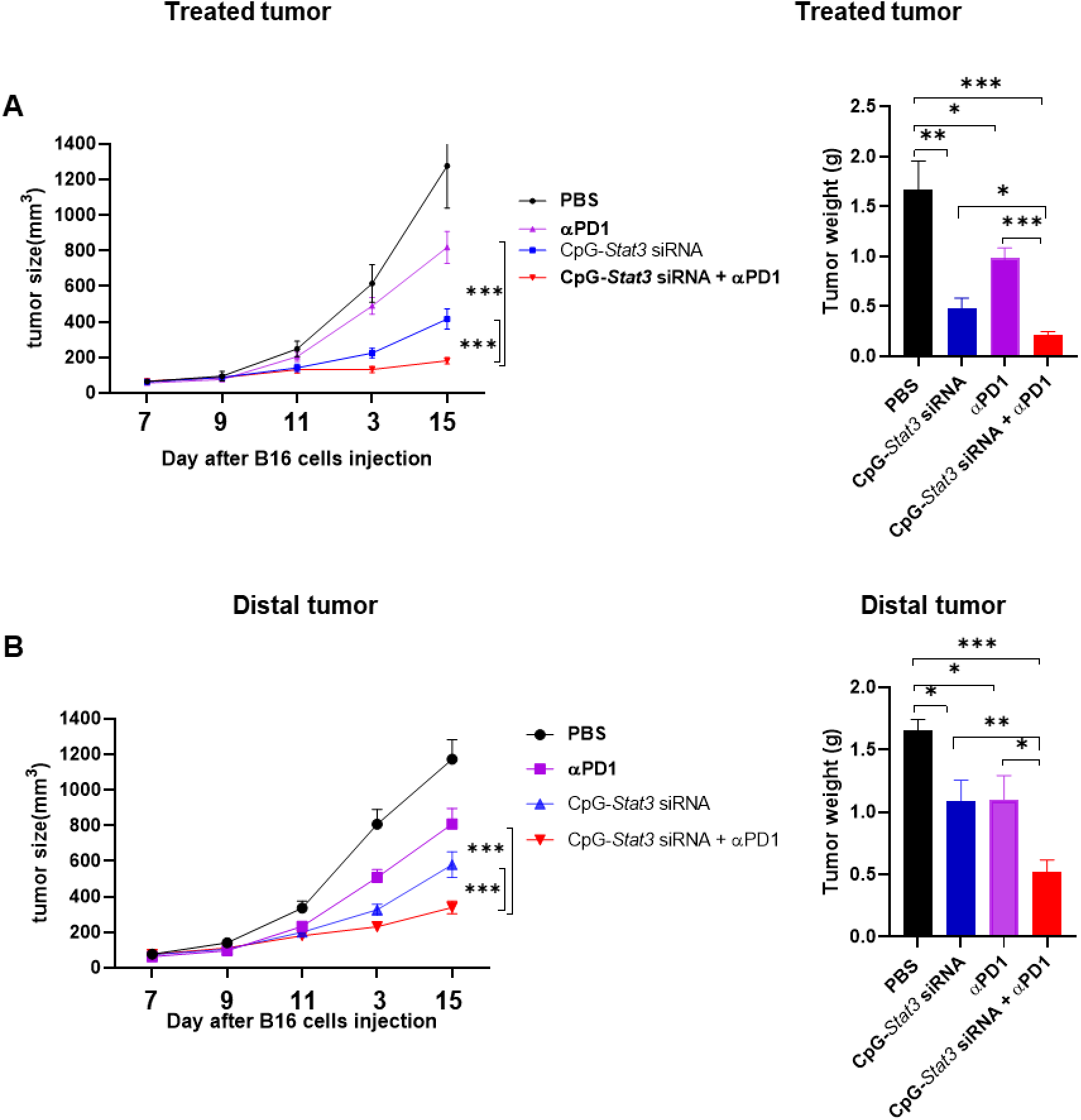
Combination treatment with intra-tumoral CpG-*Stat3* siRNA and systemic PD-1 antibody reduces growth of both treated and distal tumors. C57BL/6 mice were injected with 2 x 10^5^ B16 melanoma cells on both right and left flanks. The left flank tumors were treated by intra-tumoral injections of CpG-*Stat3* siRNA, i.p injection of anti-PD-1 antibody, or CpG-*Stat3* siRNA and anti-PD-1 every other day, starting day 7 post tumor challenge (2 x 10^5^ B16-F10 cells/tumor), n=7-8. Tumor size was monitored every other day. Data are shown as means ± SEM, Student’s *t*-test was used for statistical analysis (\**P*<0.05; \*\**P*<0.01; \*\*\**P*<0.001).

### Combination of the local CpG-*Stat3* siRNA treatment with PD-1 blockade promotes tumor T cell recruitment and activity

To assess the effect of CpG-*Stat3* siRNA/anti-PD1 combination treatment on T cell tumor infiltration and activation, we performed experiments using bilateral B16 tumor as described above. On day 15 after tumor implantation, we harvested the tumor tissues and prepared single-cell suspension, followed by antibody staining and flow cytometric analysis. Compared to PBS control, local CpG-*Stat3* siRNA treatment enhanced tumor infiltration of CD8^+^ T cells **(Figure 4A).** Intra-tumoral ratios of CD8^+^ T cells versus Tregs in treated tumor were significantly increased **(Figure 4B)**. Consistent with the antitumor effects of each treatment in **Figure 3**, local CpG-*Stat3* siRNA or systemic PD-1 antibody improved CD8^+^ T cell activity with increased IFNγ and GZMB producing CD8^+^ T cell frequency, but the most significant enhancement of the activated CD8^+^ T cells in bilateral tumors was observed in mice given the combinatory treatment **(Figure 4C)**. These results indicated that combination treatment with local CpG-*Stat3* siRNA and systemic PD-1 antibody increases tumor infiltration of activated T cells.

**Figure 4.**
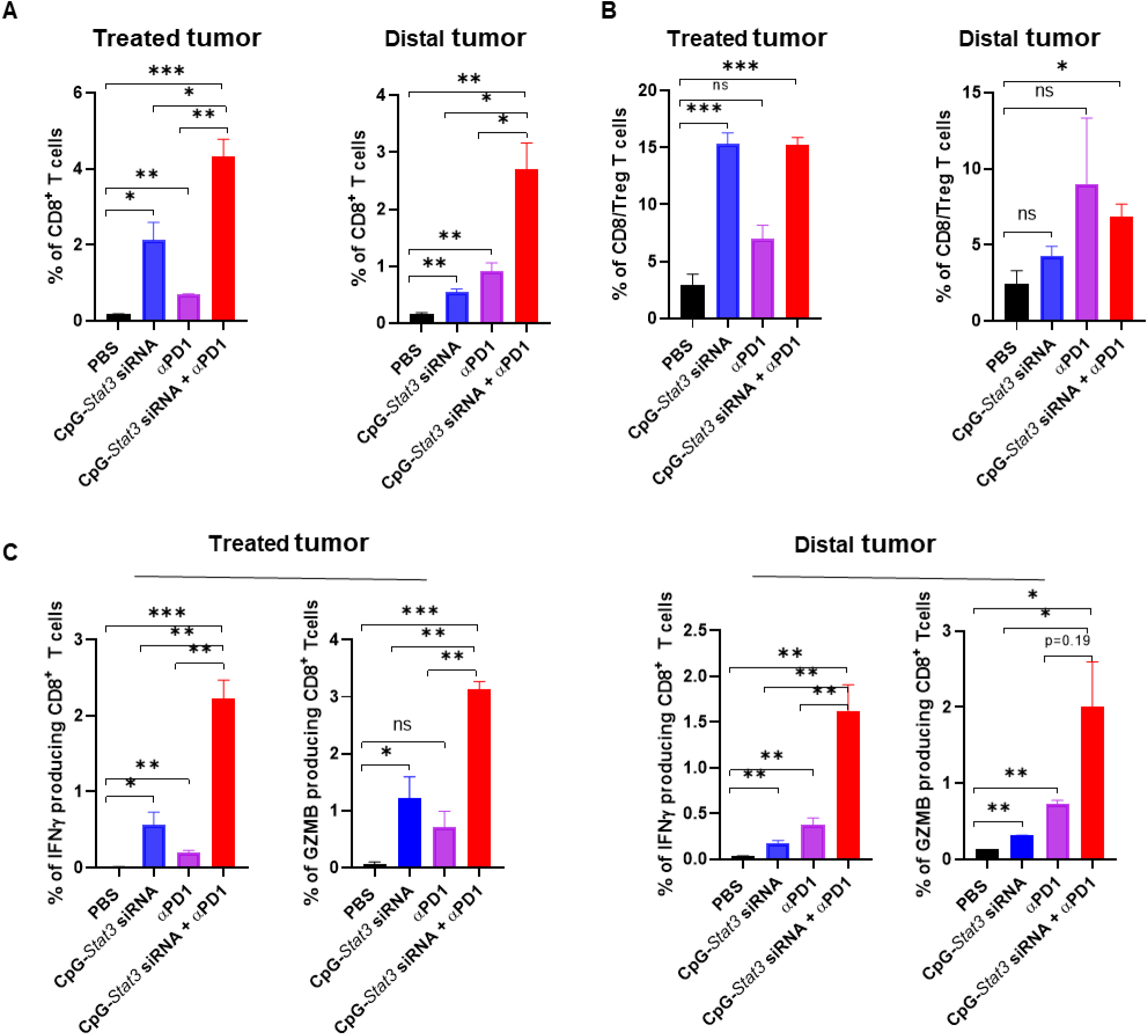
Intra-tumoral CpG-*Stat3* siRNA and systemic PD-1 antibody combined treatment significantly increases CD8^+^ T cell tumor infiltration and activity in treated and distal tumors. Single-cell suspensions from the *B16* tumors from mice with indicated treatments were analyzed by flow cytometry for CD8^+^(A) and CD8^+^/CD4^+^FoxP3^+^ immune cells(B) as well as IFNγ^+^ or GzmB^+^ CD8^+^ (C) cells from indicated treatments. Data were shown as means ± SEM, n=2-3 (n is for number of samples, each of which was pooled from 2-3 mice. Student’s t-test was used for statistical analysis (*P<0.05; **P<0.01; ***P<0.001).

### Local CpG-*Stat3* siRNA treatment systematically increases IFN**γ** and granzyme B production

In addition to immune cells, cytokines also play a critical role in anti-tumor immune therapy. Both IFNγ and GZMB are well-known for their direct cytotoxic effects on tumor cells and being able to stimulate production of immune-stimulating cytokines^42–44^. Thus, we co-cultured splenic cells from B16 tumor bearing mice receiving various treatments with B16 tumor cells. 3 or 4 days after co-culturing, the levels of IFNγ and GZMB in the cultured cell supernatants were measured by ELISA. Both IFNγ and GZMB were noticeably increased, particularly in the combinatory treatment group at day 4 after co-culturing, although there was also a significant induction of IFNγ and GZMB in local CpG-Stat3 siRNA treatment group **(Supplemental Figure S1).** These findings suggest that local CpG-Stat3 siRNA treatment induces and enhances systemic IFNγ and GZMB production triggered by PD-1 antibody treatment.

## DISCUSSION

Despite the promising outcomes of anti-CTLA4 and anti-PD-1 antibodies in treating cancer, including solid tumors and also hematopoietic malignancies^1, 3, 6, 45^, many of the treated patients do not respond well to the ICB, and those who initially do may not have durable clinical benefits^1–3^. Thus, there are urgent needs to develop immunotherapeutic strategies to improve the efficacy of ICB. The limited responses to ICB in cancer patients, including B cell lymphoma and melanoma, are largely attributed to lack of IFNγ production required for activation of CD8^+^ T cells^4–7^. Extensive studies from our group have demonstrated that STAT3, which is persistently activated in multiple types of immune cells in the tumor microenvironment, contributes to the suppression of IFNγ production in the tumor microenvironment^10, 20, 23^.

In this study, we evaluated the antitumor effects of local CpG*-Stat3* siRNA treatment at tumor sites to boost the antitumor effects of CTLA4 and/or PD-1 antibody systemic treatments in the A20 mouse lymphoma model and the B16 melanoma model. Using primary tumors as an in-situ vaccination site has been well documented^36, 46, 47^. This approach has several merits, including easier delivery, reduced drug dosing and associated toxicity, and having the potential to stimulate systemic antitumor immune responses. Our results demonstrate that combining CpG*-Stat3* siRNA with CTLA4 or PD-1 antibody suppressed A20 lymphoma tumor growth more effectively than either CpG*-Stat3* siRNA or CTLA4 or PD-1 antibody alone. In the case of B16 melanoma tumor model, we showed that CpG*-Stat3* siRNA local treatment significantly potentiates the antitumor effects of PD-1 antibody, not only at the tumors that have received the treatment, but also non-treated distal tumors. The antitumor effects on the distal non-treated tumors provide evidence supporting the activation of systemic antitumor immune responses. The detection of increased tumor CD8^+^ T cell infiltrates that are activated and able to produce IFNγ and granzyme B, and reduced tumor-associated Tregs when CpG*-Stat3* siRNA is added to ICB therapy, shows that CpG*-Stat3* siRNA can serve as an effective adjuvant for ICB. The finding that CpG*-Stat3* siRNA can elevate IFNγ and granzyme B production by splenic cells in mice receiving PD-1 further supports that CpG*-Stat3* siRNA can boost the antitumor efficacy of PD-1.

Pioneering studies by others have demonstrated CpG as an immunostimulatory molecule that activates innate immunity and induces systemic immune response through TLR9 in solid and blood cancer including B cell lymphoma and melanoma^37, 47–50^. However, high CpG dosing (30-100 μg/intratumorally) are required to enhance anti-tumor immune response in these studies^37, 47–49^. It has been demonstrated that high dose CpG injections can cause strong side effects associated with innate immune cell activation or toxicities related to chemical modification of the oligonucleotide, such as phosphorothioation^39, 40, 50^. CpG in CpG*-Stat3* siRNA acts not only as an immune stimulatory molecule but also a carrier for *Stat3* siRNA. In the present study, we showed that local injection of relatively low dose (10-15μg) of CpG-*Stat3* siRNA, in which CpG represents roughly a quarter of the conjugate, is effective in increasing both CTLA4 and PD-1 antibodies-mediated antitumor immune responses, including recruitment and activation of tumor CD8^+^ T cells, reduction of tumor infiltrating CD4^+^ Tregs and improved antitumor effects. *Stat3* gene silencing by *Stat3* siRNA in the CpG*-Stat3* siRNA conjugate has been shown to reduce the expression of immunosuppressive factors while increasing expression of immunostimulatory molecules including IFNγ^20, 29^. Because of this, it has been shown that CpG*-Stat3* siRNA is more effective in activating antitumor immune responses than CpG stimulation alone^20^. In addition to CpG, immune checkpoint antibodies are associated with autoimmune toxicities^51^. In our present study, local injection of low dose CpG-*Stat3* siRNA with reduced systemic dose PD-1 antibody (100 μg vs.200 μg) treatment can induce potent local immune modulation, resulting in systemic antitumor immune responses.

Tumor cells of both melanoma and many types of B cell lymphomas require persistently activated STAT3 for growth and/or survival^25, 26, 52, 53^. Our previous studies have shown that TLR9 is required for processing of CpG*-Stat3* siRNA to be effective siRNA^54^. B cell lymphoma cells, including mouse A20 tumor cells, express TLR9^25, 26^. Follow-up studies are needed to demonstrate the contribution of *Stat3* silencing in the tumor cells in enhancing the efficacies of ICB in A20 tumor model. Several tumor cells, especially cancer stem cells, in some solid tumors, including glioma, express TLR9^28^. The glioma cancer stem cells can also process CpG*-Stat3* siRNA into effective siRNA^28^. Nevertheless, the B16 melanoma cells do not display TLR9^20^. Therefore, at least in the B16 tumor model, CpG*-Stat3* siRNA-induced enhancement of the antitumor effects of PD-1 is largely contributed by immune activation.

Taken together, our study showed that local administration of low dose CpG-*Stat3* siRNA can enhance the therapeutic potential of IBC in B cell lymphoma and melanoma mouse tumor models, supporting further testing and development of CpG-*Stat3* siRNA and ICB combinatory treatment for clinical application.

## METHODS

### Cell lines

Mouse lymphoma cell line A20 [American Type Culture Collection (ATCC); A-20] was cultured in RPMI containing 10% fetal bovine serum (FBS: Omega Scientific) and 1 x antibiotic antimycotic (AA; Gibco), supplemented with 0.05 mM 2-mercaptoethanol (2-ME; Gibco). Mouse melanoma cell line B16 (ATCC) was cultured in Dulbeccòs modified Eaglès medium (DMEM) containing 10% FBS and 1 x AA.

### In vivo mouse experiments

Mouse care and experimental procedures were performed under pathogen-free conditions in accordance with established institutional guidance and approved protocols from Institutional Animal Care and Use Committee at Beckman Research Institute of City of Hope Medical Center. C57BL/6 and BALB/c mice were obtained from Jackson Laboratory. For the A20 lymphoma mouse model, we injected 5 x 10^5^ of A20 tumor cells subcutaneously into BALB/c mice. When the tumors reached an average size of approximately 100-150 mm^3^ (9-10 days after A20 tumor cells injection), mice with similar average tumor size were randomly divided into four groups. Then we treated A20 tumor-bearing mice with intra-tumor (IT) injections of CpG-*Stat3* siRNA (10μg/mouse), intraperitoneal (i.p) injections of CTLA4 / PD-1 antibody (100μg/mouse) or combination treatment with CpG-*Stat3* siRNA and CTLA4 / PD-1 antibody every other day. Tumor growth was monitored every other day with caliper measurement. For B16 melanoma mouse model, C57BL/6 mice were subcutaneously injected with 2 x 10^5^ of B16 tumor cells on the right and left flanks. The left flank tumors were treated by intra-tumoral injections of CpG-*STAT3* siRNA (10μg/mouse), i.p injection of PD-1 antibody (100μg/mouse), or combination treatment with CpG-*STAT3* siRNA and αPD1 every other day, starting 7 day after challenge with 2 x 10^5^ B16 cells. Tumor growth was monitored as described above.

### Oligonucleotide design and synthesis

The sequences of mouse cell-specific CpG1668(B)-siRNAs have been described previously^20^. Sequences of single-stranded constructs are listed below.

*Mouse Stat3 siRNA (SS):*

5′ CAGGGUGUCAGAUCACAUGGGCUAA 3′

*CpG1668-mouse Stat3 siRNA (AS):*

5′ TCCATGACGTTCCTGATGCT-linker-UUAGCCCAUGUGAUCUGACACCCUGAA 3′

### Quantitative Real-Time PCR

Total RNAs from various cell populations were purified with the RNeasy system according to the manufacturer’s instructions (QIAGEN). RNA (0.5 to 1 μg) was reverse transcribed to cDNA using iScript cDNA Synthesis Kit (Bio-Rad), and real-time PCR reactions were performed as described previously^10^. Specific primers for mouse Stat3, Actin and Gapdh were purchased from SA Bioscience and Qiagen. Each primer set was validated using a standard curve across the dynamic range of interest with a single melting peak. Samples were run in triplicate and expressed as means ± SEM.

### Singel cell suspension preparation

To prepare single-cell suspensions, tumor tissue was dissected into approximately 1– 5 mm^3^ fragments and digested with collagenase Type D (2 mg/ml; Roche) and DNase I (1 mg/ml; Roche) for 30–45 min at 37°C. Digests were filtered through 70 μm cell strainers, centrifuged at 1,500 rpm for 5 min. Single-cell suspensions from spleens were prepared as mentioned above. After red blood cell lysis (Sigma-Aldrich), single-cell suspensions were filtered, washed, and resuspended in FACS Buffer (2% FBS in Hank’s balanced salt solution without Ca^+^, Mg^+^, and phenol red).

### Intracellular Staining and Flow Cytometry

Intracellular staining and flow cytometric analysis were performed as previously described^10^. Single-cell suspensions (some of which were pooled from tumors harvested from 2-3 mice) were stimulated for 5 h with PMA (5 ng/ml, Sigma) and ionomycin (500 ng/ml, Sigma) in the presence of protein transport inhibitor (monensin 1000x, Biolegend). Cells were blocked with anti-CD16/CD32 antibody (TruStain FcX™ PLUS (anti-mouse CD16/32) Antibody) and incubated for 15 min on ice with PECy7-, Alexa Fluor-700, Pacific Blue (or v450), and APC-Cy7 (or Alexa Fluor-e780)-conjugated antibodies (1:100, CD4, CD8) purchased from Biolegend. After cell surface marker staining, cells were fixed and permeabilized using the BD Cytofix/Cytoperm Fixation/Permeabilization Solution Kit (BD) for IFNγ and GZMB staining or eBioscience™ Foxp3 / Transcription Factor Staining Buffer Set (eBioscience) for FoxP3 staining. Cells were then incubated with FITC, and PE-conjugated antibodies (1:100, IFNr or FoxP3, GZMB) purchased from Biolegend. Aqua LIVE/DEAD used for cell viability was purchased from Invitrogen. Cells were washed twice before analysis on the BD LSR Fortessa flow cytometer (Beckman Coulter Genomics).

### Cytokine Measurement

Splenic cells (1 × 10^5^) from B16 tumor-bearing mice were co-cultured with B16 tumor cells (2 x 10^4^) in a well of the 96-well plate for 3 or 4 days. Culture medium was collected to determine IFN-γ and GZMB levels by ELISA (R&D).

## Statistical Analysis

Statistical analyses were performed using GraphPad Prism 7 software. Statistical comparisons between groups were performed using the unpaired Student’s t test to calculate two-tailed p-value. Statistical significance values were set as ∗p<0.05. ∗∗p<0.01. ∗∗∗p<0.001. A p-value less than 0.05 would be considered statistically significant, ns = not significant. Data are presented as mean ± standard error of the mean (SEM). p value and n can be found in main and supplementary figure legends.

## DATA AVAILABILITY STATEMENT

All reagents and data generated from this study are available from the corresponding author upon a reasonable request.

## AUTHOR CONTRIBUTIONS

H.Y. and C.Z. developed the concept, designed experiments, prepared and wrote the manuscript. C.Z provided guidance and carried out the experiments, the statistical analyses. R.H. contributed to animal experiments. L.R. performed animal experiments, Q-PCR and flow cytometry experiments. J.S. performed animal experiments and Q-PCR assay. P.S. synthesized CpG-Stat3 siRNA. S. F. provided clinical insight on B cell lymphoma treatment, and M.K. contributed to the manuscript writing and discussion of the project.

## CONFILICT-OF-INTEREST STATEMENT

MK and HY are on the Scientific Advisory Board of Scopus Biopharma Inc., a licensee of the CpG-STAT3siRNA technology, with stock options. CZ also received stocks through CpG-STAT3siRNA technology licensing. MK is a scientific advisor to Duet Biotherapeutics Inc. All other authors declare no competing financial interests.

## ACKNOWLEDGMENTS

We thank the dedication of staff members at the animal facilities at City of Hope. We also acknowledge the contribution of staff members at the Analytical Cytometry Core and DNA/RNA Synthesis Core. This work is supported by the National Cancer Institute of the National Institutes of Health under grant numbers P50CA107399 (S.F.) and P30CA033572 (COH). The content is solely the responsibility of the authors and does not necessarily represent the official views of the National Institutes of Health. This study was also supported by the Billy and Audrey Wilder Endowment to H.Y.

## REFERENCES

1. Huang AC, Zappasodi R. A decade of checkpoint blockade immunotherapy in melanoma: understanding the molecular basis for immune sensitivity and resistance. Nat Immunol. 2022;23(5):660–70. Epub 2022/03/05. doi: 10.1038/s41590-022-01141-1. PubMed PMID: 35241833; PMCID: PMC9106900.

2. Morad G, Helmink BA, Sharma P, Wargo JA. Hallmarks of response, resistance, and toxicity to immune checkpoint blockade. Cell. 2021;184(21):5309–37. Epub 2021/10/09 06:00. doi: 10.1016/j.cell.2021.09.020. PubMed PMID: 34624224; PMCID: 8767569.

3. Perdikis-Prati S, Sheikh S, Bouroumeau A, Lang N. Efficacy of Immune Checkpoint Blockade and Biomarkers of Response in Lymphoma: A Narrative Review. Biomedicines. 2023;11(6). Epub 2023/06/28. doi: 10.3390/biomedicines11061720. PubMed PMID: 37371815; PMCID: PMC10296250.

4. Chen PL, Roh W, Reuben A, Cooper ZA, Spencer CN, Prieto PA, Miller JP, Bassett RL, Gopalakrishnan V, Wani K, De Macedo MP, Austin-Breneman JL, Jiang H, Chang Q, Reddy SM, Chen WS, Tetzlaff MT, Broaddus RJ, Davies MA, Gershenwald JE, Haydu L, Lazar AJ, Patel SP, Hwu P, Hwu WJ, Diab A, Glitza IC, Woodman SE, Vence LM, Wistuba, II, Amaria RN, Kwong LN, Prieto V, Davis RE, Ma W, Overwijk WW, Sharpe AH, Hu J, Futreal PA, Blando J, Sharma P, Allison JP, Chin L, Wargo JA. Analysis of Immune Signatures in Longitudinal Tumor Samples Yields Insight into Biomarkers of Response and Mechanisms of Resistance to Immune Checkpoint Blockade. Cancer discovery. 2016;6(8):827–37. Epub 2016/06/16 06:00. doi: 10.1158/2159-8290.CD-15-1545. PubMed PMID: 27301722; PMCID: 5082984.

5. Kamphorst AO, Wieland A, Nasti T, Yang S, Zhang R, Barber DL, Konieczny BT, Daugherty CZ, Koenig L, Yu K, Sica GL, Sharpe AH, Freeman GJ, Blazar BR, Turka LA, Owonikoko TK, Pillai RN, Ramalingam SS, Araki K, Ahmed R. Rescue of exhausted CD8 T cells by PD-1-targeted therapies is CD28-dependent. Science. 2017;355(6332):1423-7. doi: 10.1126/science.aaf0683. PubMed PMID: 28280249.

6. Armengol M, Santos JC, Fernandez-Serrano M, Profitos-Peleja N, Ribeiro ML, Roue G. Immune-Checkpoint Inhibitors in B-Cell Lymphoma. Cancers (Basel). 2021;13(2). Epub 2021/01/13 06:00. doi: 10.3390/cancers13020214. PubMed PMID: 33430146; PMCID: 7827333.

7. Lim SY, Shklovskaya E, Lee JH, Pedersen B, Stewart A, Ming Z, Irvine M, Shivalingam B, Saw RPM, Menzies AM, Carlino MS, Scolyer RA, Long GV, Rizos H. The molecular and functional landscape of resistance to immune checkpoint blockade in melanoma. Nature communications. 2023;14(1):1516. Epub 2023/03/20 06:00. doi: 10.1038/s41467-023-36979-y. PubMed PMID: 36934113; PMCID: 10024679.

8. Ayers M, Lunceford J, Nebozhyn M, Murphy E, Loboda A, Kaufman DR, Albright A, Cheng JD, Kang SP, Shankaran V, Piha-Paul SA, Yearley J, Seiwert TY, Ribas A, McClanahan TK. IFN-gamma-related mRNA profile predicts clinical response to PD-1 blockade. The Journal of clinical investigation. 2017;127(8):2930–40. Epub 2017/06/27 06:00. doi: 10.1172/JCI91190. PubMed PMID: 28650338; PMCID: 5531419.

9. Grasso CS, Tsoi J, Onyshchenko M, Abril-Rodriguez G, Ross-Macdonald P, Wind-Rotolo M, Champhekar A, Medina E, Torrejon DY, Shin DS, Tran P, Kim YJ, Puig-Saus C, Campbell K, Vega-Crespo A, Quist M, Martignier C, Luke JJ, Wolchok JD, Johnson DB, Chmielowski B, Hodi FS, Bhatia S, Sharfman W, Urba WJ, Slingluff CL, Jr., Diab A, Haanen J, Algarra SM, Pardoll DM, Anagnostou V, Topalian SL, Velculescu VE, Speiser DE, Kalbasi A, Ribas A. Conserved Interferon-gamma Signaling Drives Clinical Response to Immune Checkpoint Blockade Therapy in Melanoma. Cancer cell. 2020;38(4):500–15 e3. Epub 2020/09/12 06:00. doi: 10.1016/j.ccell.2020.08.005. PubMed PMID: 32916126; PMCID: 7872287.

10. Zhang C, Yue C, Herrmann A, Song J, Egelston C, Wang T, Zhang Z, Li W, Lee H, Aftabizadeh M, Li YJ, Lee PP, Forman S, Somlo G, Chu P, Kruper L, Mortimer J, Hoon DSB, Huang W, Priceman S, Yu H. STAT3 Activation-Induced Fatty Acid Oxidation in CD8(+) T Effector Cells Is Critical for Obesity-Promoted Breast Tumor Growth. Cell metabolism. 2020;31(1):148–61 e5. Epub 2019/11/26 06:00. doi: 10.1016/j.cmet.2019.10.013. PubMed PMID: 31761565; PMCID: 6949402.

11. Herrmann A, Priceman SJ, Swiderski P, Kujawski M, Xin H, Cherryholmes GA, Zhang W, Zhang C, Lahtz C, Kowolik C, Forman SJ, Kortylewski M, Yu H. CTLA4 aptamer delivers STAT3 siRNA to tumor-associated and malignant T cells. The Journal of clinical investigation. 2014;124(7):2977–87. doi: 10.1172/JCI73174. PubMed PMID: 24892807; PMCID: 4071407.

12. Yu H, Kortylewski M, Pardoll D. Crosstalk between cancer and immune cells: role of STAT3 in the tumour microenvironment. Nature reviews Immunology. 2007;7(1):41–51. doi: 10.1038/nri1995. PubMed PMID: 17186030.

13. Kortylewski M, Kujawski M, Wang T, Wei S, Zhang S, Pilon-Thomas S, Niu G, Kay H, Mule J, Kerr WG, Jove R, Pardoll D, Yu H. Inhibiting Stat3 signaling in the hematopoietic system elicits multicomponent antitumor immunity. Nature medicine. 2005;11(12):1314–21. doi: 10.1038/nm1325. PubMed PMID: 16288283.

14. Kujawski M, Kortylewski M, Lee H, Herrmann A, Kay H, Yu H. Stat3 mediates myeloid cell-dependent tumor angiogenesis in mice. The Journal of clinical investigation. 2008;118(10):3367–77. doi: 10.1172/JCI35213. PubMed PMID: 18776941; PMCID: 2528912.

15. Wang T, Niu G, Kortylewski M, Burdelya L, Shain K, Zhang S, Bhattacharya R, Gabrilovich D, Heller R, Coppola D, Dalton W, Jove R, Pardoll D, Yu H. Regulation of the innate and adaptive immune responses by Stat-3 signaling in tumor cells. Nature medicine. 2004;10(1):48–54. doi: 10.1038/nm976. PubMed PMID: 14702634.

16. Yu H, Jove R. The STATs of cancer--new molecular targets come of age. Nature reviews Cancer. 2004;4(2):97–105. doi: 10.1038/nrc1275. PubMed PMID: 14964307.

17. Deng J, Liu Y, Lee H, Herrmann A, Zhang W, Zhang C, Shen S, Priceman SJ, Kujawski M, Pal SK, Raubitschek A, Hoon DS, Forman S, Figlin RA, Liu J, Jove R, Yu H. S1PR1-STAT3 signaling is crucial for myeloid cell colonization at future metastatic sites. Cancer cell. 2012;21(5):642–54. doi: 10.1016/j.ccr.2012.03.039. PubMed PMID: 22624714; PMCID: 3360884.

18. Kortylewski M, Xin H, Kujawski M, Lee H, Liu Y, Harris T, Drake C, Pardoll D, Yu H. Regulation of the IL-23 and IL-12 balance by Stat3 signaling in the tumor microenvironment. Cancer cell. 2009;15(2):114–23. doi: 10.1016/j.ccr.2008.12.018. PubMed PMID: 19185846; PMCID: 2673504.

19. Lee H, Deng J, Kujawski M, Yang C, Liu Y, Herrmann A, Kortylewski M, Horne D, Somlo G, Forman S, Jove R, Yu H. STAT3-induced S1PR1 expression is crucial for persistent STAT3 activation in tumors. Nature medicine. 2010;16(12):1421–8. doi: 10.1038/nm.2250. PubMed PMID: 21102457; PMCID: 3088498.

20. Kortylewski M, Swiderski P, Herrmann A, Wang L, Kowolik C, Kujawski M, Lee H, Scuto A, Liu Y, Yang C, Deng J, Soifer HS, Raubitschek A, Forman S, Rossi JJ, Pardoll DM, Jove R, Yu H. In vivo delivery of siRNA to immune cells by conjugation to a TLR9 agonist enhances antitumor immune responses. Nature biotechnology. 2009;27(10):925–32. doi: 10.1038/nbt.1564. PubMed PMID: 19749770; PMCID: 2846721.

21. Lee H, Herrmann A, Deng JH, Kujawski M, Niu G, Li Z, Forman S, Jove R, Pardoll DM, Yu H. Persistently activated Stat3 maintains constitutive NF-kappaB activity in tumors. Cancer cell. 2009;15(4):283–93. doi: 10.1016/j.ccr.2009.02.015. PubMed PMID: 19345327; PMCID: 2777654.

22. Priceman SJ, Kujawski M, Shen S, Cherryholmes GA, Lee H, Zhang C, Kruper L, Mortimer J, Jove R, Riggs AD, Yu H. Regulation of adipose tissue T cell subsets by Stat3 is crucial for diet-induced obesity and insulin resistance. Proceedings of the National Academy of Sciences of the United States of America. 2013;110(32):13079–84. doi: 10.1073/pnas.1311557110. PubMed PMID: 23878227; PMCID: 3740863.

23. Zhang C, Xin H, Zhang W, Yazaki PJ, Zhang Z, Le K, Li W, Lee H, Kwak L, Forman S, Jove R, Yu H. CD5 Binds to Interleukin-6 and Induces a Feed-Forward Loop with the Transcription Factor STAT3 in B Cells to Promote Cancer. Immunity. 2016;44(4):913–23. Epub 2016/04/21 06:00. doi: 10.1016/j.immuni.2016.04.003. PubMed PMID: 27096320; PMCID: 4844010.

24. Scuto A, Kujawski M, Kowolik C, Krymskaya L, Wang L, Weiss LM, Digiusto D, Yu H, Forman S, Jove R. STAT3 inhibition is a therapeutic strategy for ABC-like diffuse large B-cell lymphoma. Cancer Res. 2011;71(9):3182–8. Epub 2011/04/28. doi: 10.1158/0008-5472.CAN-10-2380. PubMed PMID: 21521803; PMCID: PMC3085657.

25. Zhang Q, Hossain DM, Nechaev S, Kozlowska A, Zhang W, Liu Y, Kowolik CM, Swiderski P, Rossi JJ, Forman S, Pal S, Bhatia R, Raubitschek A, Yu H, Kortylewski M. TLR9-mediated siRNA delivery for targeting of normal and malignant human hematopoietic cells in vivo. Blood. 2013;121(8):1304–15. Epub 2013/01/05. doi: 10.1182/blood-2012-07-442590. PubMed PMID: 23287859; PMCID: PMC3578952.

26. Zhao X, Zhang Z, Moreira D, Su YL, Won H, Adamus T, Dong Z, Liang Y, Yin HH, Swiderski P, Pillai RK, Kwak L, Forman S, Kortylewski M. B Cell Lymphoma Immunotherapy Using TLR9-Targeted Oligonucleotide STAT3 Inhibitors. Mol Ther. 2018;26(3):695–707. Epub 2018/02/13. doi: 10.1016/j.ymthe.2018.01.007. PubMed PMID: 29433938; PMCID: PMC5910676.

27. Li X, Wei Y, Wei X. Napabucasin, a novel inhibitor of STAT3, inhibits growth and synergises with doxorubicin in diffuse large B-cell lymphoma. Cancer Lett. 2020;491:146–61. Epub 2020/08/18. doi: 10.1016/j.canlet.2020.07.032. PubMed PMID: 32798587.

28. Herrmann A, Cherryholmes G, Schroeder A, Phallen J, Alizadeh D, Xin H, Wang T, Lee H, Lahtz C, Swiderski P, Armstrong B, Kowolik C, Gallia GL, Lim M, Brown C, Badie B, Forman S, Kortylewski M, Jove R, Yu H. TLR9 is critical for glioma stem cell maintenance and targeting. Cancer Res. 2014;74(18):5218–28. Epub 2014/07/23 06:00. doi: 10.1158/0008-5472.CAN-14-1151. PubMed PMID: 25047528; PMCID: 4167470.

29. Herrmann A, Kortylewski M, Kujawski M, Zhang C, Reckamp K, Armstrong B, Wang L, Kowolik C, Deng J, Figlin R, Yu H. Targeting Stat3 in the myeloid compartment drastically improves the in vivo antitumor functions of adoptively transferred T cells. Cancer Res. 2010;70(19):7455–64. Epub 2010/09/16 06:00. doi: 10.1158/0008-5472.CAN-10-0736. PubMed PMID: 20841481; PMCID: 3058618.

30. Kortylewski M, Kujawski M, Herrmann A, Yang C, Wang L, Liu Y, Salcedo R, Yu H. Toll-like receptor 9 activation of signal transducer and activator of transcription 3 constrains its agonist-based immunotherapy. Cancer Res. 2009;69(6):2497–505. Epub 2009/03/05 09:00. doi: 10.1158/0008-5472.CAN-08-3031. PubMed PMID: 19258507; PMCID: 2657819.

31. Herrmann A, Lahtz C, Nagao T, Song JY, Chan WC, Lee H, Yue C, Look T, Mulfarth R, Li W, Jenkins K, Williams J, Budde LE, Forman S, Kwak L, Blankenstein T, Yu H. CTLA4 Promotes Tyk2-STAT3-Dependent B-cell Oncogenicity. Cancer Res. 2017;77(18):5118–28. Epub 2017/07/19 06:00. doi: 10.1158/0008-5472.CAN-16-0342. PubMed PMID: 28716895; PMCID: 5600851.

32. Garris CS, Arlauckas SP, Kohler RH, Trefny MP, Garren S, Piot C, Engblom C, Pfirschke C, Siwicki M, Gungabeesoon J, Freeman GJ, Warren SE, Ong S, Browning E, Twitty CG, Pierce RH, Le MH, Algazi AP, Daud AI, Pai SI, Zippelius A, Weissleder R, Pittet MJ. Successful Anti-PD-1 Cancer Immunotherapy Requires T Cell-Dendritic Cell Crosstalk Involving the Cytokines IFN-gamma and IL-12. Immunity. 2018;49(6):1148–61 e7. Epub 2018/12/16 06:00. doi: 10.1016/j.immuni.2018.09.024. PubMed PMID: 30552023; PMCID: 6301092.

33. Hsu FJ, Komarovskaya M. CTLA4 blockade maximizes antitumor T-cell activation by dendritic cells presenting idiotype protein or opsonized anti-CD20 antibody-coated lymphoma cells. Journal of immunotherapy (Hagerstown, Md: 1997). 2002;25(6):455–68. Epub 2002/11/20 04:00. doi: 10.1097/00002371-200211000-00002. PubMed PMID: 12439343.

34. Kepp O, Marabelle A, Zitvogel L, Kroemer G. Oncolysis without viruses – inducing systemic anticancer immune responses with local therapies. Nature reviews Clinical oncology. 2020;17(1):49–64. Epub 2019/10/09 06:00. doi: 10.1038/s41571-019-0272-7. PubMed PMID: 31595049.

35. Broomfield SA, van der Most RG, Prosser AC, Mahendran S, Tovey MG, Smyth MJ, Robinson BW, Currie AJ. Locally administered TLR7 agonists drive systemic antitumor immune responses that are enhanced by anti-CD40 immunotherapy. Journal of immunology (Baltimore, Md: 1950). 2009;182(9):5217–24. Epub 2009/04/22 09:00. doi: 10.4049/jimmunol.0803826. PubMed PMID: 19380767.

36. Oba T, Makino K, Kajihara R, Yokoi T, Araki R, Abe M, Minderman H, Chang AE, Odunsi K, Ito F. In situ delivery of iPSC-derived dendritic cells with local radiotherapy generates systemic antitumor immunity and potentiates PD-L1 blockade in preclinical poorly immunogenic tumor models. Journal for immunotherapy of cancer. 2021;9(5). Epub 2021/05/30 06:00. doi: 10.1136/jitc-2021-002432. PubMed PMID: 34049930; PMCID: 8166607.

37. Reilley MJ, Morrow B, Ager CR, Liu A, Hong DS, Curran MA. TLR9 activation cooperates with T cell checkpoint blockade to regress poorly immunogenic melanoma. Journal for immunotherapy of cancer. 2019;7(1):323. Epub 2019/11/28 06:00. doi: 10.1186/s40425-019-0811-x. PubMed PMID: 31771649; PMCID: 6880482.

38. Gao C, Kozlowska A, Nechaev S, Li H, Zhang Q, Hossain DM, Kowolik CM, Chu P, Swiderski P, Diamond DJ, Pal SK, Raubitschek A, Kortylewski M. TLR9 signaling in the tumor microenvironment initiates cancer recurrence after radiotherapy. Cancer Res. 2013;73(24):7211–21. Epub 2013/10/25 06:00. doi: 10.1158/0008-5472.CAN-13-1314. PubMed PMID: 24154870; PMCID: 3917779.

39. Schmidt U, Wagner H, Miethke T. CpG-DNA upregulates the major acute-phase proteins SAA and SAP. Cellular microbiology. 1999;1(1):61–7. Epub 2001/02/24 11:00. doi: 10.1046/j.1462-5822.1999.00007.x. PubMed PMID: 11207541.

40. von Beust BR, Johansen P, Smith KA, Bot A, Storni T, Kundig TM. Improving the therapeutic index of CpG oligodeoxynucleotides by intralymphatic administration. European journal of immunology. 2005;35(6):1869–76. Epub 2005/05/24 09:00. doi: 10.1002/eji.200526124. PubMed PMID: 15909311.

41. van der Leun AM, Thommen DS, Schumacher TN. CD8(+) T cell states in human cancer: insights from single-cell analysis. Nature reviews Cancer. 2020;20(4):218–32. Epub 2020/02/07 06:00. doi: 10.1038/s41568-019-0235-4. PubMed PMID: 32024970; PMCID: 7115982.

42. Velotti F, Barchetta I, Cimini FA, Cavallo MG. Granzyme B in Inflammatory Diseases: Apoptosis, Inflammation, Extracellular Matrix Remodeling, Epithelial-to-Mesenchymal Transition and Fibrosis. Frontiers in immunology. 2020;11:587581. Epub 2020/12/03 06:00. doi: 10.3389/fimmu.2020.587581. PubMed PMID: 33262766; PMCID: 7686573.

43. Wensink AC, Hack CE, Bovenschen N. Granzymes regulate proinflammatory cytokine responses. Journal of immunology (Baltimore, Md: 1950). 2015;194(2):491–7. Epub 2015/01/04 06:00. doi: 10.4049/jimmunol.1401214. PubMed PMID: 25556251.

44. Boulch M, Cazaux M, Cuffel A, Guerin MV, Garcia Z, Alonso R, Lemaitre F, Beer A, Corre B, Menger L, Grandjean CL, Morin F, Thieblemont C, Caillat-Zucman S, Bousso P. Tumor-intrinsic sensitivity to the pro-apoptotic effects of IFN-gamma is a major determinant of CD4(+) CAR T-cell antitumor activity. Nature cancer. 2023;4(7):968–83. Epub 2023/05/30 01:06. doi: 10.1038/s43018-023-00570-7. PubMed PMID: 37248395; PMCID: 10368531.

45. Goodman A, Patel SP, Kurzrock R. PD-1-PD-L1 immune-checkpoint blockade in B-cell lymphomas. Nature reviews Clinical oncology. 2017;14(4):203–20. Epub 2016/11/03 06:00. doi: 10.1038/nrclinonc.2016.168. PubMed PMID: 27805626.

46. Hammerich L, Binder A, Brody JD. In situ vaccination: Cancer immunotherapy both personalized and off-the-shelf. Molecular oncology. 2015;9(10):1966–81. Epub 2015/12/04 06:00. doi: 10.1016/j.molonc.2015.10.016. PubMed PMID: 26632446; PMCID: 5528727.

47. Proskurina AS, Ruzanova VS, Ritter GS, Efremov YR, Mustafin ZS, Lashin SA, Burakova EA, Fokina AA, Zatsepin TS, Stetsenko DA, Leplina OY, Ostanin AA, Chernykh ER, Bogachev SS. Antitumor efficacy of multi-target in situ vaccinations with CpG oligodeoxynucleotides, anti-OX40, anti-PD1 antibodies, and aptamers. Journal of biomedical research. 2022;37(3):194–212. Epub 2023/05/10 12:42. doi: 10.7555/JBR.36.20220052. PubMed PMID: 37161885; PMCID: 10226083.

48. Sagiv-Barfi I, Czerwinski DK, Levy S, Alam IS, Mayer AT, Gambhir SS, Levy R. Eradication of spontaneous malignancy by local immunotherapy. Science translational medicine. 2018;10(426). Epub 2018/02/02 06:00. doi: 10.1126/scitranslmed.aan4488. PubMed PMID: 29386357; PMCID: 5997264.

49. Sagiv-Barfi I, Kohrt HE, Burckhardt L, Czerwinski DK, Levy R. Ibrutinib enhances the antitumor immune response induced by intratumoral injection of a TLR9 ligand in mouse lymphoma. Blood. 2015;125(13):2079–86. Epub 2015/02/11 06:00. doi: 10.1182/blood-2014-08-593137. PubMed PMID: 25662332; PMCID: 4375105.

50. Krieg AM. CpG still rocks! Update on an accidental drug. Nucleic acid therapeutics. 2012;22(2):77–89. Epub 2012/02/23 06:00. doi: 10.1089/nat.2012.0340. PubMed PMID: 22352814.

51. Tahir SA, Gao J, Miura Y, Blando J, Tidwell RSS, Zhao H, Subudhi SK, Tawbi H, Keung E, Wargo J, Allison JP, Sharma P. Autoimmune antibodies correlate with immune checkpoint therapy-induced toxicities. Proceedings of the National Academy of Sciences of the United States of America. 2019;116(44):22246–51. Epub 2019/10/16 06:00. doi: 10.1073/pnas.1908079116. PubMed PMID: 31611368; PMCID: 6825284.

52. Fu XQ, Liu B, Wang YP, Li JK, Zhu PL, Li T, Tse KW, Chou JY, Yin CL, Bai JX, Liu YX, Chen YJ, Yu ZL. Activation of STAT3 is a key event in TLR4 signaling-mediated melanoma progression. Cell death & disease. 2020;11(4):246. Epub 2020/04/22 06:00. doi: 10.1038/s41419-020-2440-1. PubMed PMID: 32312954; PMCID: 7171093.

53. Liu YX, Xu BW, Niu XD, Chen YJ, Fu XQ, Wang XQ, Yin CL, Chou JY, Li JK, Wu JY, Bai JX, Wu Y, Li SM, Yu ZL. Inhibition of Src/STAT3 signaling-mediated angiogenesis is involved in the anti-melanoma effects of dioscin. Pharmacological research. 2022;175:105983. Epub 2021/11/26 06:00. doi: 10.1016/j.phrs.2021.105983. PubMed PMID: 34822972.

54. Nechaev S, Gao C, Moreira D, Swiderski P, Jozwiak A, Kowolik CM, Zhou J, Armstrong B, Raubitschek A, Rossi JJ, Kortylewski M. Intracellular processing of immunostimulatory CpG-siRNA: Toll-like receptor 9 facilitates siRNA dicing and endosomal escape. Journal of controlled release: official journal of the Controlled Release Society. 2013;170(3):307–15. Epub 2013/06/20 06:00. doi: 10.1016/j.jconrel.2013.06.007. PubMed PMID: 23777886; PMCID: 3742675.

