## Supplemental Figure for "Local CpG-*Stat3* siRNA treatment improves antitumor effects of immune checkpoint inhibitors"

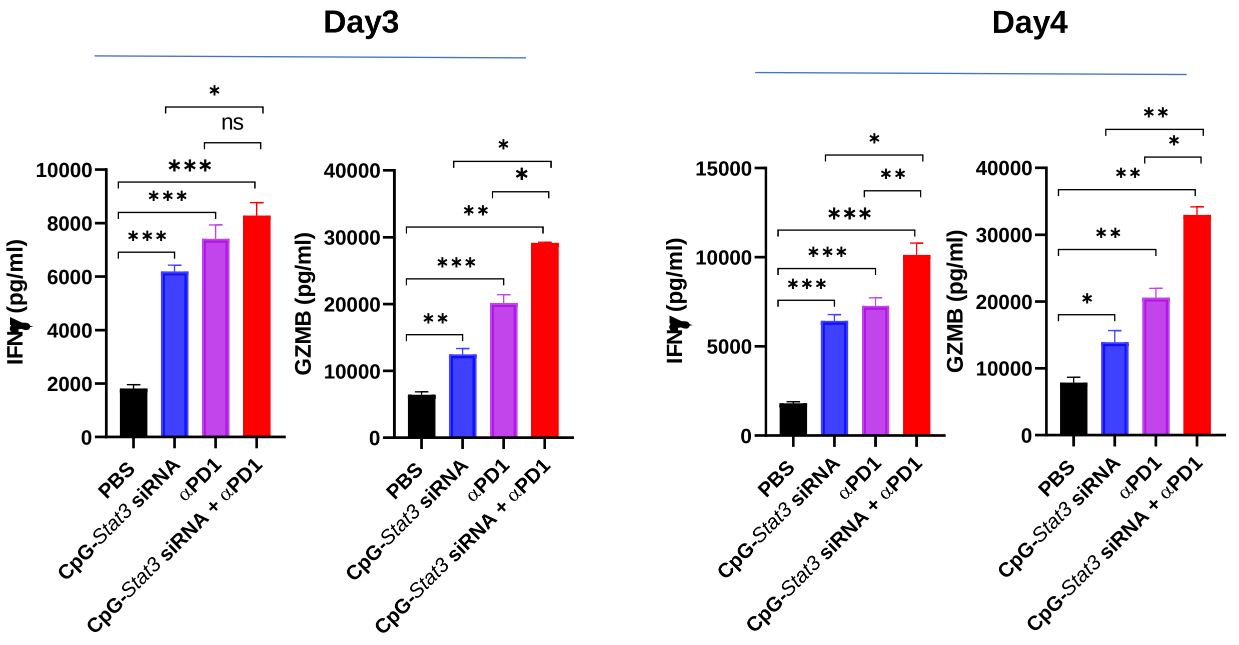


**Supplemental Figure S1 : Systematic IFN-γ production and GZMB expression after intra-tumoral CpG-*Stat3* siRNA and systemic PD-1 antibody single or combined treatments.** Splenic cells from B16 tumor bearing mice with different treatment were co-cultured with B16 tumor cells for 3 or 4 days. The supernatants from co-culture cells were collected. ELISA assays of the co-cultured supernatant were performed to quantify secreted IFNγ and Granzyme B, both of which are the main cytotoxic molecules released by effector T cells. Data were shown with means ± SEM, n=2-3 (n is for number of samples, each of which was pooled from 2-4 mice. Student`s *t*-test was used for statistical analysis (*P<0.05; **P<0.01; ***P<0.001).
